# Back to basics: Observed statistics are sufficient to predict drug responses

**DOI:** 10.64898/2026.06.09.731197

**Authors:** Valentine Svensson, Umair Khan, Hamed Heydari, Airol A. Ubas, Nicole Thomas, Daniele Merico, Hani Goodarzi, John Yu, Nima Alidoust, Shreshth Gandhi

## Abstract

Predicting how cells, tissues, and patients will respond to a drug, cytokine, or genetic perturbation is central to biological and clinical reasoning. The practical goal is to estimate the analysis-ready readouts that support this reasoning: which cellular responses are context-dependent, which perturbations reveal shared or divergent mechanisms, and which observations should become the basis for the next experiment.

Here we introduce Rhaister, a perturbation-response predictor that operates directly on screen-level summary statistics. By measuring just a few perturbations in a new biological context, Rhaister predicts the unmeasured perturbations by learning how response patterns vary across reference contexts. This formulation applies to both fine-grained molecular readouts, such as transcriptional responses from Tahoe-100M or other large perturbation screen, or on phenotypic endpoints. To train and apply Rhaister on phenotypic endpoints we created Emerald Bay, a purpose-built Tahoe dataset that unifies multi-day cancer drug perturbation, pooled Mosaic tumor contexts, and paired transcriptomic response measurements^*^.

Across these settings, Rhaister matches or exceeds substantially more expensive virtual-cell models, often achieving the highest values possible in evaluation metrics, while training in seconds and running predictions in milliseconds. On Emerald Bay, Rhaister predicts context-specific drug phenotypes from sensitivity measurements alone and improves further when including transcriptomic information. We also introduce Rhaister-O, predicting drug responses in new contexts from baseline expression alone and, to our knowledge, provides the first zero-shot model for this task. Rhaister establishes summary-statistic perturbation modeling as a fast, interpretable framework for predicting biological response across new contexts.

## Introduction

Predicting how a sample (a cell population, a tissue, a patient) responds to a perturbation sits at the core of biological reasoning. Nominating targets, finding selective perturbations, identifying which patients they should reach, and reasoning through how a perturbation acts and how resistance emerges all leverage this prediction among others.

In practice we do this in two steps. We measure how perturbations act in disease models that are accessible and scalable enough to reason through, and we use that to predict what happens in contexts we cannot reach as easily.

Both steps are bounded. Running perturbations in accessible models is feasible but slow. It ranges from weeks to months per screen, and for longer across the tens or hundreds of thousands of perturbations we would want to probe. As biological data and simulators become coupled to agents that reason in seconds, that latency becomes disqualifying. The contexts that matter most are harder still: rare cell types, tissues that will not grow outside the body, diseased cells that resist culture, and cells inside patients reachable only through clinical trials. Even if time wouldn’t be a limiting factor, perturbing these at scale would be impossible.

For broad value in reasoning, we need diverse readouts. Often we want the response of specific pathways; just as often we want direct phenotypic endpoints that matter for discovery, e.g., cell death across cancer cells of many genetic backgrounds.

Together, these define the problem: infer diverse phenotypes from perturbations in new biological samples.

There have been many attempts using various innovative “virtual cell models” for this purpose [1], [2], [3], [4], [5], [6], [7]. These models are used to simulate single-cell data of perturbations in new contexts. Three gaps remain: zero-shot prediction, reaching more complex phenotypes, and pace of training / prediction for integration of these models as grounding data in agentic reasoning [8], [9]. Furthermore, no model has shown clear scaling with more perturbations of samples.

Here we present our modeling framework “Rhaister”, designed to predict transcriptional perturbation responses and other phenotypes. Our process inverts the typical single-cell perturbation modeling workflow (Figure 1, top right). Rhaister operates on aggregated summary statistics directly from large and diverse data to enable training and prediction at the speed of agentic reasoning systems.

**Figure 1.**
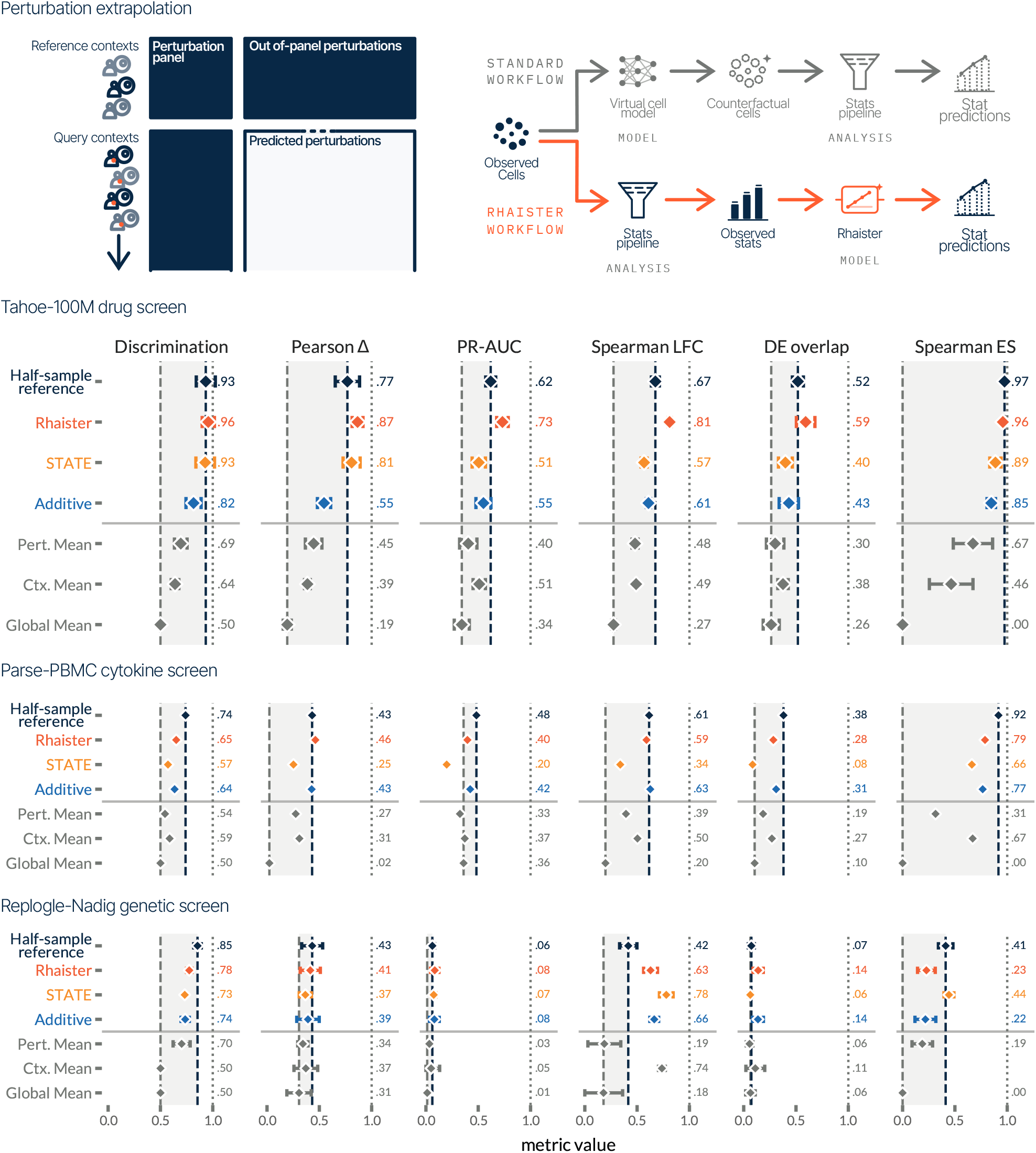
**Perturbation extrapolation, top left:** Schematic of the perturbation extrapolation task. Our goal is to predict how a new biological context will respond to *any* drug in our library given how that context responds to a small panel of drugs. **Top right**: Schematic of our modeling workflow, contrasted against typical perturbation prediction workflows. **Tahoe-100M drug screen:** Results on Tahoe-100M using six evaluation metrics reflecting downstream analysis tasks. Confidence intervals represent ± 1SD across train/test splits. **Parse-PBMC cytokine screen, Replogle-Nadig genetic screen:** Performance on additional perturbation screen data using the same evaluation metrics.

To enable modeling of phenotypic readout, we created Emerald Bay, a new perturbation atlas of cancer therapeutics applied to a large number of biological contexts, providing drug phenotype readout with paired transcriptional readout from two million single cells. Using this data, we apply Rhaister to the task of drug phenotype prediction, where it substantially outperforms baselines.

Using only summary statistics, the Rhaister model achieves state of the art performance in general across evaluation metrics, and in many cases saturating performance. Rhaisters performance is in large part enabled by rich and diverse training data. Structurally, Rhaister learns to predict perturbation responses as combinations of panel perturbations. With a large and sufficiently diverse training set, this straightforward strategy is sufficient to saturate predictive performance using a small panel of observed drugs.

Finally, having achieved maximal possible performance when predicting perturbation responses from a drug panel using Rhaister, we moved to the more ambitious task of “zeroshot” prediction. We developed “RhaisterO”, a variation on Rhaister predicting perturbation responses using baseline transcriptomic features alone. This would enable translation of perturbation effects from accessible in vitro models to inaccessible arbitrary biological contexts.

Rhaister-O is the first model to meaningfully improve over simple baseline models for the baseline-only perturbation prediction task, and outperforming complex prior state of the art virtual cell models for this task.

By changing the focus from accurately simulating single cells for the sake of downstream analysis, to modeling analysis results, we can make more effective use of large perturbation screens. Screens with high diversity and rich readouts like *Tahoe-100M* and *Emerald Bay* provide strong predictive signals when focusing on the effects of perturbations.

### Rhaister saturates transcriptional perturbation prediction evaluation metric performance

Transcriptional perturbation responses of gene regulation are reflected in three units that are output from statistical analysis of single-cell gene expression data: log2 fold change, Mann-Whitney *U*-test p-values, and expression delta. These summary statistics are estimated by comparing a perturbation to a corresponding negative control for each biological context. They are the typical output artifacts a researcher uses to interpret the results of a perturbation, reflecting which genes’ expressions are affected by the perturbation, how strongly, and to what degree of certainty.

The summary statistics for each combination of (context, perturbation, gene) are treated as different views of the data, and our goal is to predict all three views, for all genes, in unseen (context, perturbation) combinations, given seen combinations of (context, perturbation).

The Rhaister model expresses the summary statistics of *non-panel perturbations* as linear combinations of summary statistics of *panel perturbations*. The linear weights are trained using *reference contexts* where the full perturbation screen of both non-panel and panel perturbations are available. The model can then be applied to *query contexts* where only panel perturbations have been observed to produce predictions of the summary statistics.

We applied the Rhaister model to Tahoe-100M drug screening data [10]. Evaluating prediction using six metrics reflecting properties important for downstream computational biology analysis [2], shows Rhaister as overcoming naive group-mean baselines and an additive baseline, as well as the state-of-the-art virtual cell model STATE (Figure 1). The Rhaister model performance is on par with half-sample reference performance, a measure of how well the observed data can *predict itself*, indicating that performance on these evaluation metrics is saturated, with no meaningful benefit to improving upon them.

The rich combinations of contexts and perturbations available in the Tahoe-100M data enable this straightforward yet successful framing: unseen perturbations as linear combinations of seen perturbations. Having access to many reference contexts allows the model to accurately learn which perturbations are informative in which contexts.

To evaluate broad applicability of Rhaister, we applied it to predict donor-specific perturbation responses in a cytokine screen on peripheral blood from human donors [11]. In the cytokine screen dataset, an additive baseline showed strong performance across the evaluation metrics. The Rhaister model slightly outperformed the additive baseline, approaching half-sample reference performance. Both the additive baseline and Rhaister outperformed the STATE model (Figure 1 Parse-PBMC cytokine screen).

Finally, we also applied the Rhaister model to a gene editing CRISPR screen dataset limited to only four biological contexts [12], [13]. In this case Rhaister achieves saturated performance on several evaluation metrics, but under-performs on other metrics relative to STATE (Figure 1 Replogle-Nadig genetic screen). This is consistent with predictive performance being enabled by having access to a large number of reference contexts for training.

### Emerald Bay enables drug phenotype prediction

In the context of cancer, the role of therapeutics is to kill cancer cells, or at least inhibit their growth. The key drug phenotype is sensitivity to drugs in these effects. Defining drug sensitivity by relative growth rate inhibition (relative decrease in cell count compared to vehicle) is frequently used as the primary phenotype. Context-specific drug sensitivity is of particular interest, since selectivity for certain cancer subtypes based on shared genetics or transcriptional states is used as a proxy for selectivity for those cancers.

To train models for drug sensitivity prediction, we created Emerald Bay, a dataset from a multi-day treatment of biologically pooled Mosaic tumors [10], [14] against a battery of cancer therapeutics. Emerald Bay contained a curated set of anticancer agents at multiple doses treated across a Mosaic tumor pool optimized for five-day culture conditions. The Mosaic tumor pool represents a diverse set of solid tumors (Figure 2 Emerald Bay cancer models). Emerald Bay data is available on HuggingFace^1^.

**Figure 2.**
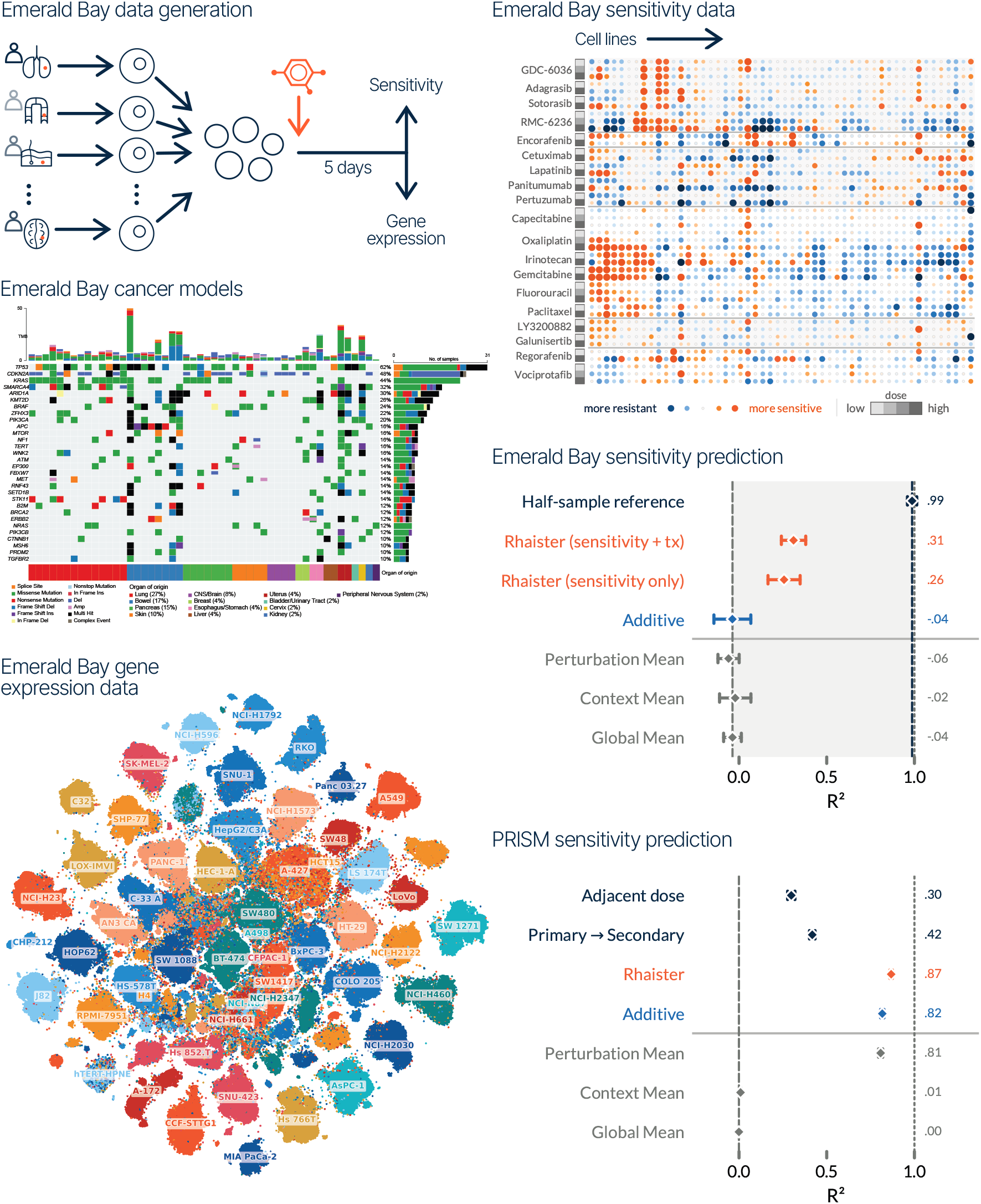
**Emerald Bay data generation:** Schematic of the Mosaic data generation process. **Emerald Bay cancer models:** Oncoplot of the cell lines showing tumor mutational burden (TMB), recurrent driver alterations, and tissue of origin. **Emerald Bay gene expression data:** TSNE visualization of the single cell gene expression data. **Emerald Bay sensitivity data:** Drug sensitivity phenotypic data from the Emerald Bay dataset. **Emerald Bay sensitivity prediction:** Predictive performance on Emerald Bay sensitivity data. Confidence intervals represent ±1SD across five train/test splits. **PRISM sensitivity prediction:** Similar for the PRISM dataset.

We applied Rhaister to Emerald Bay, to predict drug sensitivity as a summary statistic per context and perturbation. In Emerald Bay, each combination of (context, perturbation) has a sensitivity summary statistic. Our goal is to learn from observed (context, perturbation) combinations with a panel of drugs, what will happen for unseen perturbations (drugs treatments) in new cell contexts. No naive baselines perform successfully (coefficient of determination, *R*^2^, of ∼0), while Rhaister predictions have *R*^2^, of 0.26 (Figure 2 Emerald Bay sensitivity prediction).

In addition to context-specific sensitivity, the Emerald Bay dataset has single-cell whole transcriptome gene expression in 1,800,000 cells. From these we can derive summary statistics of transcriptomic changes due to the drug perturbations serving as additional predictors of drug sensitivity. When Including transcriptional response summary statistics^2^ from the drug panel as additional predictors, Rhaister performance increases to 0.31 *R*^2^.

Rhaister also outperforms naive baselines on the PRISM drug sensitivity screen data, with an *R*^2^ of 0.87. In the PRISM sensitivity data, the perturbation mean baseline show high performance (*R*^2^ of 0.81), with better predictive performance on the secondary screen than the primary screen results (*R*^2^ of 0.42) or the adjacent dose of the same drug (*R*^2^ of 0.30). This likely reflects the focus on broadly effective drugs in this dataset, without specificity to biological contexts. Several drugs are either systematically ineffective across contexts or systematically effective across contexts (especially chemotherapeutics at high dose).

Together, these results highlight Emerald Bay as a uniquely enabling dataset for learning context-specific drug sensitivity from paired perturbational phenotypes and transcriptional responses. By supporting multi-day drug perturbations across diverse pooled cancer contexts, the Mosaic platform provides a scalable experimental framework for evaluating and improving predictive models of therapeutic response such as Rhaister. Prediction of drug sensitivity is important for therapeutic pipelines, and we are making the Emerald Bay dataset publicly available for the community to train models and evaluate models with.

### Performance of perturbation prediction scales with the perturbation panel size

The number of panel drugs needed to predict a full drug library is a key design factor governing the scalability and economics of collecting data. Smaller panels need less material simplifies the logistics of data collection.

Predictive information available for the Rhaister model scales directly with the number of panel perturbations used for prediction. The Rhaister model expresses each non-panel perturbation as a linear combination of panel perturbations. With larger panels, more types of perturbation responses can be represented.

To quantify the information gained by increasing the number of panel perturbations, we subsampled the number of drugs (each at three doses) available in the observed panel to use for predictions of the non-panel perturbations in the Tahoe-100M dataset. Across all evaluation metrics, we do not reach diminishing returns on increasing the size of the panel (Figure 3 Left). On average, halving the number of drugs in the panel reduced the performance by ∼1.5%. The number of perturbations to use in a panel should be balanced between desired performance and available budget.

**Figure 3.**
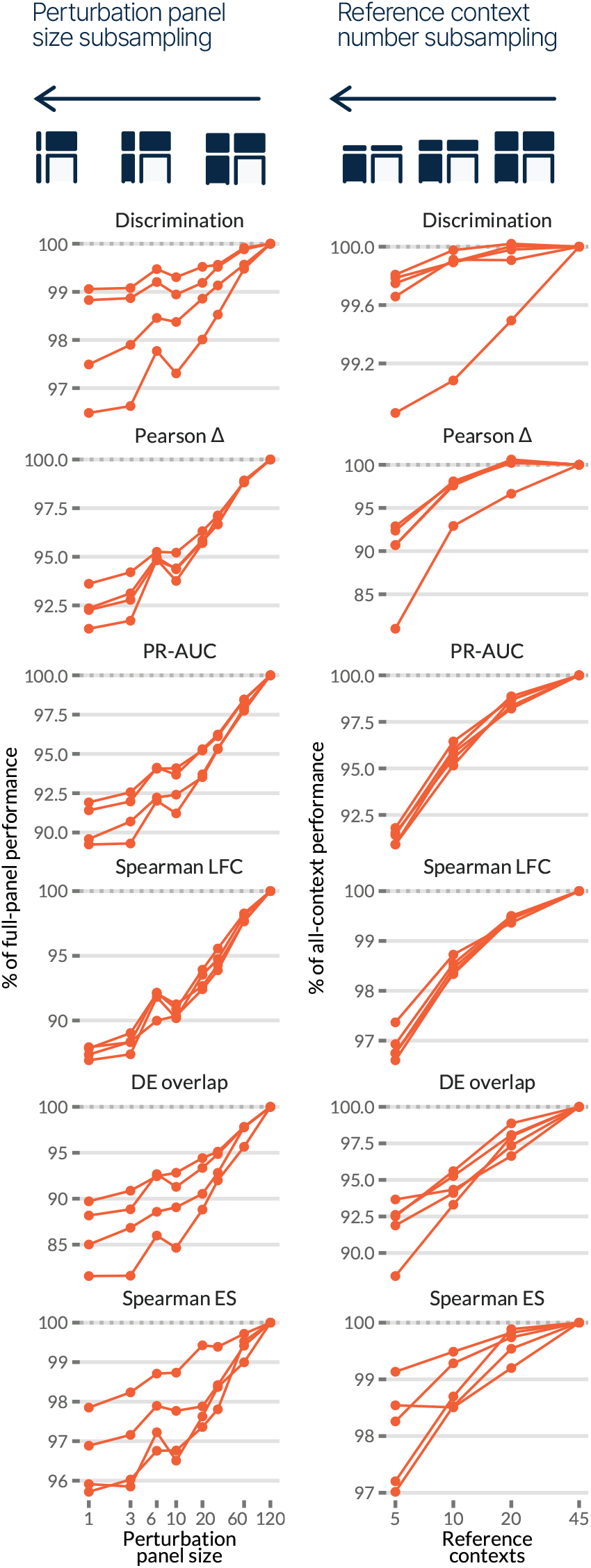
**Left:** Relative improvement in performance metrics with increasing numbers of unique drug responses used as predictors. **Right:** Relative improvement in performance with increasing numbers of reference contexts used for training.

This is in contrast with results from subsampling the number of reference contexts used for training. We subsampled the number of cell lines available as reference during training. The relative increase in performance is marginal after ∼20 reference contexts are available (Figure 3 Right).

Even with a small panel of a single drug at three doses, perturbation prediction performance reach the level of the half-sample reference performance across the metrics. This predictive performance is reached using 10 cell lines as reference contexts.

### Rhaister outperforms virtual cell modeling in predicting perturbation from basal state data alone

An ultimate and ambitious goal for perturbation modeling is to predict perturbation responses given *only* basal state gene expression. This would enable prediction of perturbation responses in vast catalogs and atlases of available baseline gene expression data.

As a separate task branch in the Rhaister modeling framework, we formulated Rhaister-O, to specifically use only information available from unperturbed controls to predict responses. We refer to this variation as ‘zeroshot’ since we have seen zero perturbation responses from a query context, and cannot make use of similarity between perturbations.

Our summary statistics-based zeroshot prediction model outperforms the state of the art virtual cell model STATE for this zeroshot prediction task across all evaluation metrics (Figure 4).

**Figure 4.**
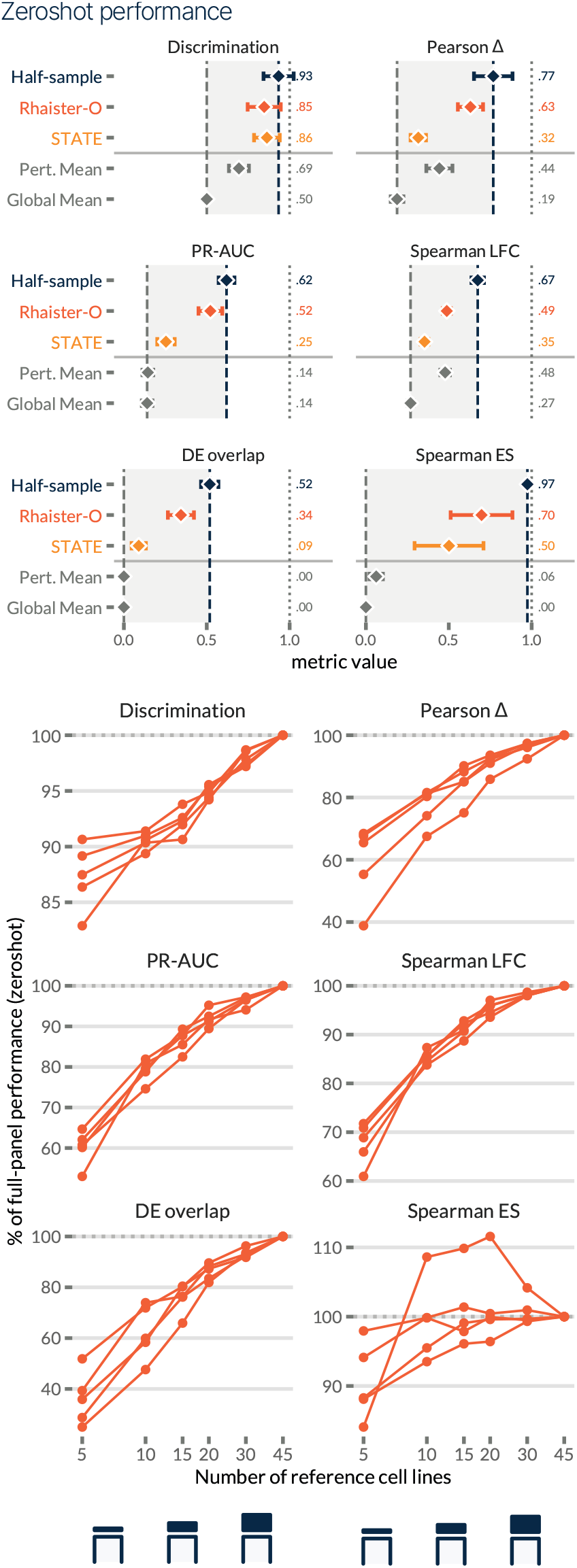
**Top:** Performance of zeroshot perturbation prediction in new contexts with only baseline expression data. Confidence intervals reflect ±1SD across five train/test splits. **Bottom:** Relative improvement in performance with increasing number reference contexts used for training.

We subsampled the number of reference context used for training of the Rhaister-O model. While the results show a pattern of diminishing returns across metrics, the metrics do not appear to have plateaued.

Increasing the number of reference contexts available for training the perturbation response prediction model is likely to improve performance in the zeroshot setting.

The gap in performance could be interpreted in how information is used by the models. In the fewshot setting, each query perturbation is interpreted as a combination of reference perturbations. While in the zeroshot setting, there is no such direct relation between predictors.

## Discussion

Reasoning models now generate hypotheses and insights from the existing literature in seconds. What they cannot generate is the novel ground truth those hypotheses must be tested against. In biology that ground truth is expensive and slow to produce, so a large corpus of high-quality data, already generated and ready to be consumed, is what keeps verification from becoming the bottleneck.

No corpus, however large or diverse, will cover every context, and the contexts that matter most are often the hardest to measure. This is our motivation: to build the datasets, such as Tahoe-100M and Emerald Bay, and, on top of them, models that generalize their signals to unseen harder-to-reach-contexts.

Virtual cell models have promised exactly this, operating on individual cells and aggregating up the readouts used for downstream analysis. Even where they succeed, they fall short on the dimensions that matter for reasoning at scale: they are slow to train and to run, they do not extend readily to direct phenotypes, and they have not delivered zero-shot prediction. At points, they have struggled to improve on simple baselines at all.

This motivated us to go back to basics. Rather than simulate cells and aggregate, we model the observed statistics directly.

We present a new modeling strategy for predicting drug treatment effects using drug panels from large scale perturbation data. A key insight is to move predictive modeling from biological primitives to summary statistics. This simplifies data management and enables straightforward models to achieve good predictive performance.

This limits the strategy from the ability to perform arbitrary single-cell level tasks based on simulated data, as is the goal for virtual cell models. It is possible to imagine scenarios where counterfactual simulation produces distributional properties of interest (emergence of previously unknown cell types, novel lineage commitment biases, unexpected synergistic gene module coexpression, etc). In practical terms however, vast amounts of perturbation responses that are important and useful for follow-up discussion occur in static, pre-defined, cellular contexts.

The Rhaister^3^ modeling framework operates directly on precisely the summary statistics that are already of interest for downstream analysis. The same summary statistics that would be of interest from a new screening data batch for a researcher are the ones that can be predicted from a panel with the Rhaister framework.

Through the Emerald Bay data, we have seen that this is not limited to transcriptional responses. While a model that works within a summary statistic view saturates performance for transcriptional drug response data, the increased performance on the sensitivity prediction task points to other cases and summary statistics of interest where information can be shared between summary statistics to improve predictive performance.

A related observation is for the Rhaister-O case of “zeroshot” prediction. It is likely possible to obtain good population-level predictors for new contexts by leveraging other predictors available from different biological contexts at baseline untreated states.

A key problem remains: expanding Rhaister to predicting response in harder to perturb biological samples, which is the subject of subsequent studies. In the meantime, Rhaister provides a framework for interpretable predictions of biological phenotypes in unseen biological samples rapidly. It transfers the bottleneck in prediction from complexity in modeling methods towards scaling data generation, with accuracies comparable to experimental variation, expanding the ground truth data available for verifying output of reasoning models in different biological tasks.

## Details

### The Rhaister model

The goal of the Rhaister model is to learn how contexts and perturbations interact to produce perturbation responses. As a starting point, we fit a noninteraction, purely additive model

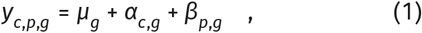

where *c* indexes contexts, *p* indexes perturbations, and *g* indexes genes. The additive model is fitted using alternating least squares.

Additive predictions for an unseen combination (*c*^⋆^, *p*^⋆^) a query context *c*^⋆^, and a non-panel perturbation *p*^⋆^, can be made by

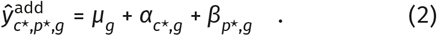

For a query context, *c*^⋆^, we form a ridge regression problem that predicts an unseen perturbation *p*^⋆^ as a linear combination of observed perturbations in *c*^⋆^. Let 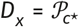 be perturbations observed for reference contexts 𝒞 _ref_. Then we stack refence context observations into 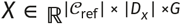, and the targets for the unseen perturbation *p*^⋆^ into 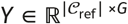, letting us solve

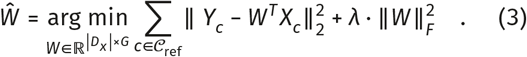

Predictions can then be made by

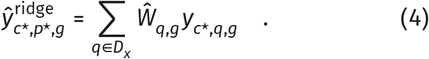

In cases where we are lacking coverage in reference data, we fall back to additive predictions:

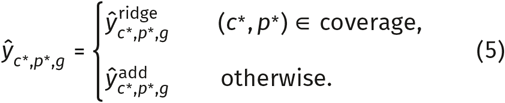

For perturbation responses, we have access to multiview data reflecting different aspects of responses: log fold change, *Y*^LFC^; Mann-Whitney U-test (MWU) p-values, *Y*^MWU^, and expression deltas, *Y*^Δ^. Specifically for the MWU view, predictions are calibrated using a small multilayer perceptron, taking as input a number of available summary statistics,

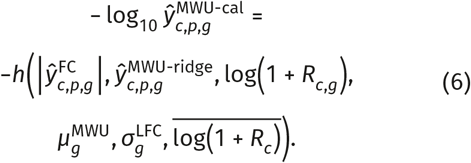

Here *R*_*c,g*_ is the baseline unperturbed gene expression of gene *g* in context *c*, 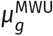 is the mean MWU p-value per gene across training samples, 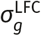 is the standard <u>deviation</u> of fold changes across training samples, and log(1 + *R*_*c*_) is averaged over genes.

Calibration predictions are evenly blended with regression-based MWU predictions:

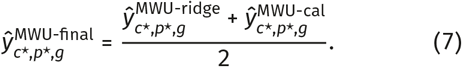

### The Rhaister-O zeroshot model

The Rhaister model requires having seen target summary statistics for a panel of perturbations in a number of reference contexts.

A more ambitious goal is to be able to predict perturbations responses from baseline expression alone. This would enable perturbation response predictions in a vast catalog of available baseline gene expression data.

The Rhaister-O variation of the modeling framework assumes we have no panel perturbations to reference, but we do instead have access to baseline expression summary statistics. Using baseline pseudobulk gene expression *x* as predictor summary statistics, we can fit a model

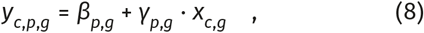

using reference contexts *c*, where all perturbations have been observed. From this model, predictions can be made through

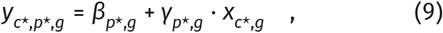

since we are assuming that we can obtain untreated control expression data 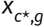 from a new (query) context 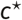.

### Baselines and reference models

Unless otherwise noted, models are trained on the training fraction of a dataset and train/test split.

#### Global mean

The global mean baseline assumes that perturbation responses look the same regardless of which context or perturbation the data is from,

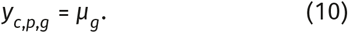

Global means fitted to the training data is used for predictions,

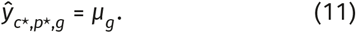

#### Context mean

The context mean baseline assumes that different biological contexts respond the same way to different perturbations, regardless of what the perturbation is,

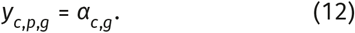

Predictions are the per-context mean responses fitted from the training data,

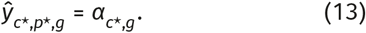

#### Perturbation mean

Analogously to context mean, the perturbation mean baseline assumes that a perturbation will have the same effect regardless of biological context of the cells it is applied to,

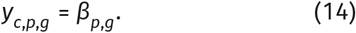

Predictions are the perperturbation mean responses fit ted from the training data,

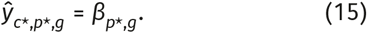

#### Additive

Assuming perturbation responses depend on both biological context and perturbation, but these effects being independent, leads to an additive baseline model,

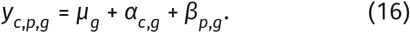

The additive baseline model is fitted through alternating least squares (ALS) on the training data, and predictions can be made for unseen (*c*^⋆^, *p*^⋆^) combinations through

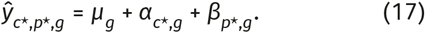

#### State Transport

State Transport (STATE) is a conditional set transport model operating on sets of single cells. Given gene expression profiles for a set of (observed) unperturbed, cells, and a conditioning signal (e.g., perturbation identity), the individual cells are transported to gene expression profiles compatible with perturbed cells.

Transported, counterfactual, cells from unobserved perturbations can subsequently be passed through the same summary statistics pipeline as cells from observed perturbations, resulting in summary statistics predictions 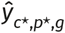.

#### Half-sample reference

Single cells from original single-cell data were divided into two equally sized parallel datasets per context, perturbation, and batch. These two datasets have the same possible groupings as the original dataset, and can be passed through the same summary statistics pipeline to produce results *Y* ^*A*^ and *Y* ^*B*^.

By evaluating ‘predictions’ 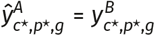 (and vice versa) for the test set of a dataset and split, we can learn approximately optimal performance for a combination of dataset, metric, and statistical pipeline.

It is worth noting that assignment of cells in to parallel sets is stochastic, and each of *Y* ^*A*^ and *Y* ^*B*^ will be the result of half as much input data as original summary statistics *Y*. This means that half-sample reference performance values are not hard ceilings on performance.

#### Permute Context & Permute Perturbation

To confirm that the Rhaister model is learning interaction signals based on the training set, we perform ablation of the information content in the training data by randomly permuting either context labels or perturbation labels in the training data. Models trained on permuted data are applied to unpermuted test data.

#### Primary → Secondary

In high throughput drug screens, such as PRISM, results from a primary screen is used to nominate followup perturbations in a secondary screen. As a baseline, we use the results of the primary screen as a prediction for the secondary screen (which we use for Rhaister modeling),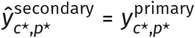.

#### Adjacent dose

As an alternative positive control baseline in the PRISM dataset, we use adjacent dose results of a matched drug as predictors for the drug, dose combination to be predicted.

### Data

The Rhaister model is applied to summary statistics from multiple perturbation datasets.

#### Tahoe-100M drug perturbation screen

The Tahoe-100M dataset is an scRNA-seq dataset of 100 million cells from 50 different cell lines having been exposed to 384 different drugs at three different concentrations [10]. Each cell line is treated as a biological context. Perturbation effects were quantified by performing comparative analysis between treatment and negative control independently for each cell line, within each plate, to ensure that batch-matched samples are compared.

We create five independent train-test splits on the context level, in each defining five cell lines a as query contexts and the remaining 45 cell lines as reference contexts. For those five query contexts, 120 drugs were defined as panel perturbations, reserving 240 drugs as non-panel perturbations. Each dosage of each of the drugs is treated as an independent perturbation in practice, leaving a total of 360 panel perturbations and 720 non-panel perturbations.

#### Parse-PBMC cytokine screen

The Parse-PBMC dataset consist of scRNA-seq data measuring the response of 90 cytokines in 18 peripheral blood cell types collected from 12 donors [11]. The combinations of donor and cell type are considered as biological contexts. We made use of a previously defined split where all cell types from four donors are held out as query contexts^4^. Of the cytokines, 28 are defined as panel-perturbations, leaving 62 non-panel perturbations. Cytokine treatment vs untreated comparisons were used to create perturbations response statistics for each donor and cell type combination independently.

#### Replogle-Nadig genetic peturbation screen

The Replogle-Nadig dataset is a combination of multiple CRISPR genetic screen datasets using a shared set of 1,700 genetic perturbations [12], [13]. The combined dataset consist of four different cell lines which we treated as biological context. Differential expression and expression deltas were estimated using non-targeting guides as negative control for each target, using cells matched by cell line and GEM group as a batch identifier. Summary statistics were averaged over batches for each cell line, gDNA target combination.

We used four previously defined train-test splits, where in each split one cell line is held out as query context using the other three cell lines as reference contexts. In each split, between 400 and 500 CRISPR targets were used as panel perturbations, while keeping 1,000 CRISPR targets as non-panel perturbations to be predicted.

The combined dataset was obtained from HuggingFace^5^.

#### Emerald Bay

A new dataset generated with the MOSAIC platform, similar to Tahoe-100M [10] but smaller-scale and customized for longduration drug treament was used for sensitivity prediction. Emerald Bay contained 91 drug treatments (including combination treatments) across 52 cell lines. Cells were harvested 5 days after drug or DMSO treatments, yielding 1,831,648 cells passing full QC filters and an extra 291,765 passing minimal QC filters. Bioinformatics processing was similar to Tahoe-100M, with fastq files first processed using split-pipe followed by genotype-based cell line demultiplexing based on Demuxlet. Cell counts used for sensitivity quantification uses the minimal QC filters, while gene expression data used for additional predictors of sensitivity uses full QC filters.

Sensitivity for each context (cell line) and perturbation combination was estimated as

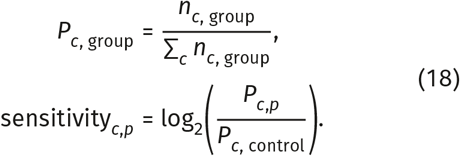

For Rhaister modeling, the dataset was filtered to 45 singled-rug perturbations, covering 19 unique drugs at 23 doses.

We created five independent train-test splits on the context level, in each defining five cell lines as query contexts and the remaining 47 cell lines as reference contexts. For those five query contexts, six unique drugs with their 2-3 different doses were defined as panel perturbations, reserving 14 unique drugs (with their 2-3 doses) as non-panel perturbations serving as targets for prediction.

Summary statistics for Emerald Bay are available on HuggingFace^6^.

#### PRISM sensitivity screen

The PRISM sensitivity screen is a dataset using a molecular barcoding method to to measure growth inhibition activity of drugs in human cancer cell lines [15].

As a summary statistic for sensitivity, we used the log2FC (log2 fold-change) as provided in the PRISM dataset^7^, which corresponds to the log treatment/DMSO ratio of the cell line abundance estimate (based on normalized Luminex median fluorescence intensity).

### Summary statistics

There are many proposed statistical pipelines to produce different summary statistics for perturbation responses. The choice of summary statistics depends on the specific properties of a perturbation of interest.

#### Log2 fold change

This statistic is meant to indicate the relative difference in expression levels between perturbed and unperturbed cells. We use log2 fold change (L2FC) for gene expression of each gene *g* as defined by the software package `pdex` ^8^, where it is estimated by

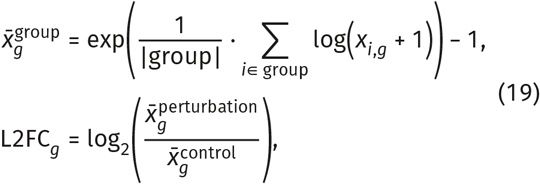

where *x*_*i,g*_ are UMI counts of gene *g* in cell *i*.

#### Mann-Whitney U-test p-value

An alternative quantification of perturbation response is through relative ordering of expression levels among cells, quantified through a Mann-Whitney U-test (MWU) from the `pdex` software package. The `pdex` package uses standard two-sided MWU with tie-correction and continuity correction. If perturbed cells are systematically ranked high or low, the null hypothesis can be rejected according to the MWU statistic, which is converted to a p-value, and further adjusted through FDR correction.

#### Expression delta

An alternative to log2 fold change is often used in single cell perturbation screens known as *expression delta*, Δ, defined in the `cell-eval` ^9^ software package as

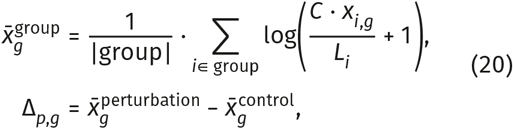

where 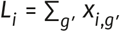 is each cell’s raw library size and *C* is a scaling constant.

#### Sensitivity

Sensitivity measures the effectivity of a drug inhibiting growth rate of a cell context. Sensitivity statistics are defined separately for the Emerald Bay and PRISM datasets, and used as-is from the input data.

### Evaluation metrics

In most cases, evaluation metrics are calculated independently for each combination (*c*^⋆^, *p*^⋆^) of query contexts and non-panel perturbations, followed by arithmetic mean averaging across the combinations.

#### Pearson correlation of Δ

This quantifies consistency of relative magnitudes between observed gene expression Deltas 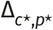 and pre-dicted gene expression Deltas 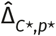,

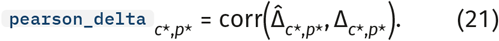

#### Perturbation discrimination

We adapt the perturbation discrimnation score [16] for the setting of perturbation effects using observed and predicted gene expression Deltas,

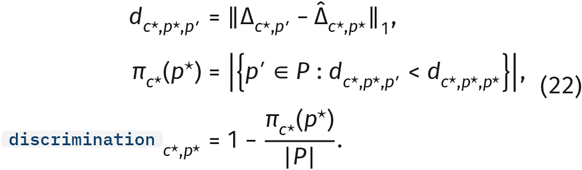

This metric reflects relative ranking of similarity from a predicted perturbation to observed perturbations. A high score indicates that even though absolute similarity to a true perturbation is poor, the prediction is more similar to the correct perturbation than other perturbations.

#### PR-AUC of differential expression classification

Differential expression classification quantifies whether predicted per-gene Mann-Whitney U-test p-values falling under a certain threshold matches true genes’ MWU p-values falling under the same threshold:

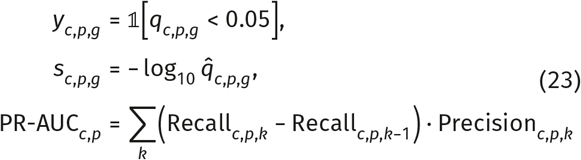

We use the threshold of 0.05 on the FDR transformed scale of the p-values.

#### Spearman LFC

The Spearman LFC metric quantifies whether genes passing an MWU p-value threshold have a consistent rank-ordering of log2 fold changes between true and predicted perturbation results,

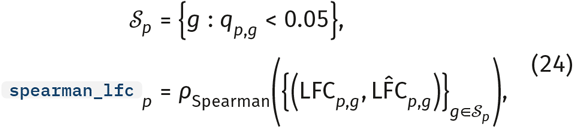

#### DE overlap

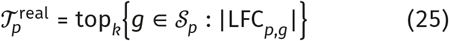

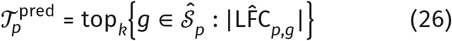

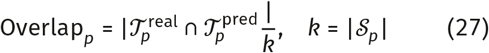

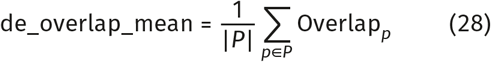

#### Spearman effect size

The authors of STATE defined transcriptome wide perturbation ‘effect size’ as the number of genes that pass a MWU p-value threshold. This metric quantifies rank ordering of effect sizes for perturbations per biological context,

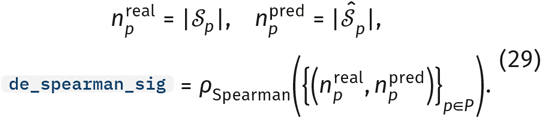

#### Coefficient of determination (*R*^2^)

The coefficient of determination *R*^2^ measures the proportion of residual variance after applying predictions over the the total variance of the observed data,

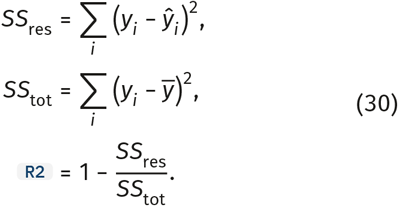

## Additional material

**Figure S1.**
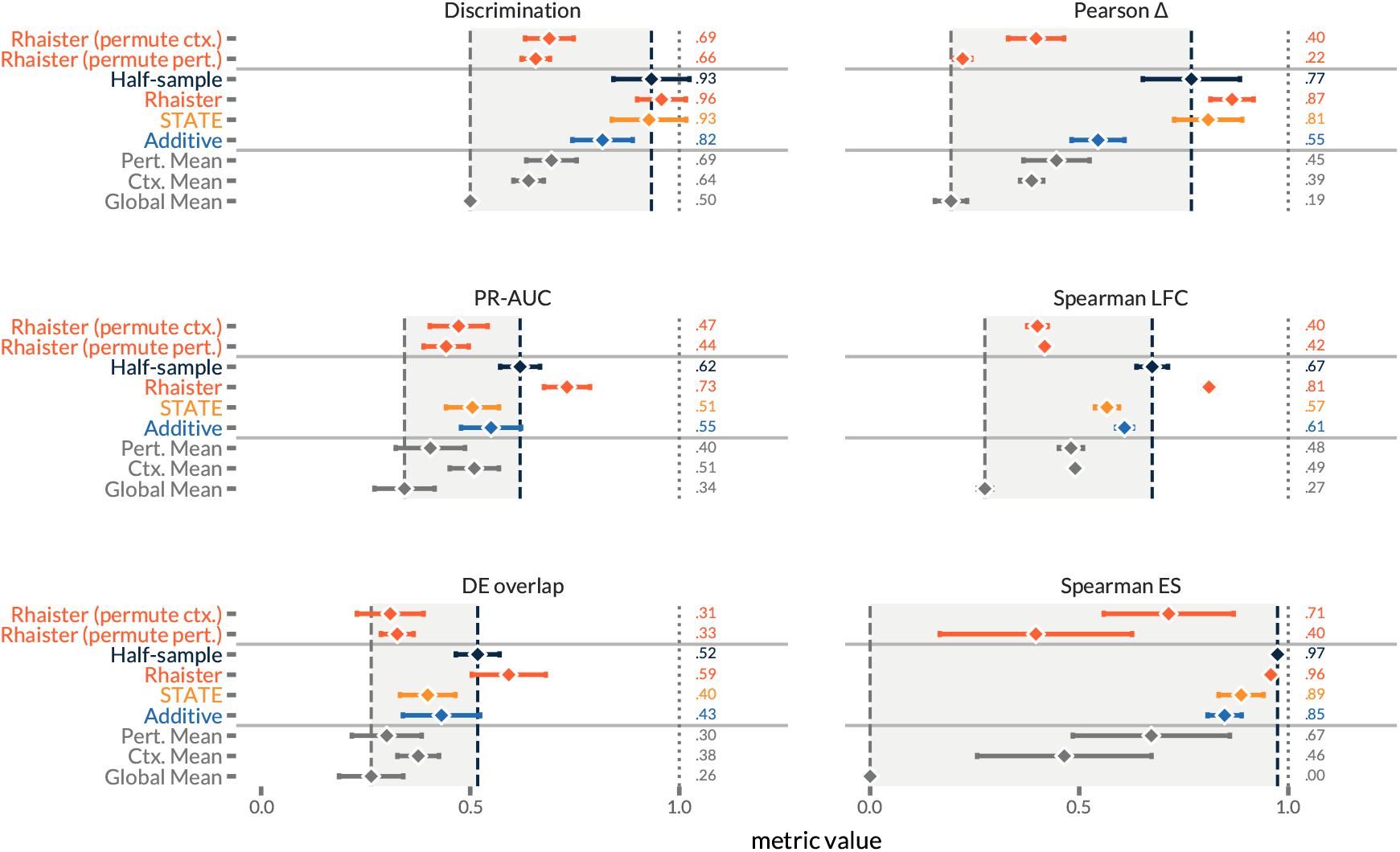
Performance comparison with training signal ablations for Rhaister.

## Footnotes

* https://huggingface.co/collections/tahoebio/rhaister

1 https://huggingface.co/datasets/tahoebio/EmeraldBay

2 expression delta, differential expression, MWU statistics

3 https://huggingface.co/tahoebio/Rhaister

4 https://github.com/ArcInstitute/state/blob/main/state_preprint_toml_files/parse_tomls/donor.toml

5 https://huggingface.co/datasets/arcinstitute/State-Replogle-Filtered

6 https://huggingface.co/datasets/tahoebio/EmeraldBay

7 https://depmap.org/repurposing/

8 https://github.com/ArcInstitute/pdex

9 https://github.com/ArcInstitute/celleval

